# Phenotypic parallelism during experimental adaptation of a free-living bacterium to the zebrafish gut

**DOI:** 10.1101/2020.03.18.997734

**Authors:** Jarrett F. Lebov, Brandon H. Schlomann, Catherine D. Robinson, Brendan J. M. Bohannan

**Affiliations:** Institute of Ecology and Evolution, University of Oregon, Eugene, Oregon, USA; Institute of Molecular Biology, University of Oregon, Eugene, Oregon, USA; Department of Physics, Materials Science Institute, University of Oregon, Eugene, Oregon, USA; Institute for Genome Sciences, University of Maryland, Baltimore, Maryland, USA

## Abstract

Despite the fact that animals encounter a plethora of bacterial species throughout their lives, only a subset are capable of colonizing vertebrate digestive tracts, and these bacteria can profoundly influence the health and development of their animal hosts. However, it is still unknown how bacteria evolve symbioses with animal hosts, and this process is central to both the assembly and function of gut bacterial communities. Therefore, we used experimental evolution to study a free-living bacterium as it adapts to a novel vertebrate host. We serially passaged replicate populations of *Shewanella oneidensis*, through the digestive tracts of larval zebrafish (*Danio rerio*). After only 20 passages, representing approximately 200 bacterial generations, isolates from replicate evolved populations displayed an improved ability to colonize larval zebrafish digestive tracts during competition against their unpassaged ancestor. Upon sequencing the genomes of these evolved isolates, we discovered that the two isolates with the highest mean competitive fitness accumulated unique sets of mutations. We characterized the swimming motility and aggregation behavior of these isolates, as these phenotypes have previously been shown to alter host-microbe interactions. Despite exhibiting different biofilm characteristics, both isolates evolved augmented swimming motility. These enhancements are consistent with expectations based on the behavior of a closely related *Shewanella* strain previously isolated from the zebrafish digestive tract and suggest that our evolved isolates are pursuing a convergent adaptive trajectory with this zebrafish isolate. In addition, parallel enhancements in swimming motility among isolates from independently adapted populations implicates increased dispersal as an important factor in facilitating the onset of host association. Our results demonstrate that free-living bacteria can rapidly improve their associations with vertebrate hosts.

## Introduction

Bacterial lineages have radiated into practically every imaginable niche on Earth [1, 2]. In particular, the vertebrate digestive tract houses bacterial communities whose composition is distinct from those found in surrounding environments [3, 4], and this suggests that host-associated bacteria maintain certain traits that enable them to colonize animal hosts. In order to establish and maintain host-association, bacteria must surmount a multitude of complex challenges, including traversing diverse physical landscapes, harvesting energy from dynamic nutrient sources, and protecting themselves from antimicrobial compounds. Thus, the number of traits involved in host-association is likely enormous [5, 6]. Despite this complexity, previous analyses indicate that novel host-microbe symbioses have arisen multiple times throughout evolutionary history [7]. However, it is unknown which suites of traits enable bacteria to transition to host association, or how likely they are to evolve.

It is well established that bacteria residing in vertebrate digestive tracts have substantial impacts on the health and development of their animal hosts [8–11]. Consequently, many researchers have sought to understand which traits provide bacteria the capacity to colonize the vertebrate gut [12–17], but this body of work has relied almost exclusively on snapshots of host-microbe relationships after they have evolved. Because information may be lost over the course of evolution, it is difficult to infer traits that facilitate transitions to host association by limiting examinations solely to strains that have already made such a transition [18].

To better understand how bacteria initiate host associations, we took an experimental evolution approach involving the serial passage of a free-living bacterial strain through the digestive tracts of a model vertebrate. When combined with genomic sequencing of evolving lineages, this strategy enables the observation of evolutionary changes in genotype at fine temporal scales [19]. Phenotypic and fitness assays can then be performed, and this information can be synthesized to understand which sets of traits facilitate adaptation, how those traits interact to improve fitness, and in what order those traits are likely to evolve [20–23].

Many insights have been gained from employing this type of approach *in vitro*, and experimental evolution practitioners have begun expanding into *in vivo* environments to investigate evolutionary dynamics within hosts [24–28]. A recent example of this was conducted by Robinson and colleagues who experimentally adapted an *Aeromonas* strain previously isolated from zebrafish (*Danio rerio*) to germ free (GF) larval zebrafish to explore how this bacterium might increase its association with its vertebrate host [29]. They found that evolved *Aeromonas* isolates achieved higher relative abundances in the digestive tracts of larvae during competition against their ancestral strain. Further, these increases in relative fitness appeared to be explained by augmented motility of evolved isolates, leading to faster rates of colonization.

For their study, Robinson et al. used a bacterial symbiont that had been isolated from a zebrafish gut, and thus it is not known how this *Aeromonas* species’ relationship with zebrafish originated, or which traits may have been involved in the initiation of this process. Therefore, while this previous work investigated how established bacterial symbionts can improve their ability to colonize the host, we sought to understand how a bacterium might initiate a novel host-microbe symbiosis. We accomplished this by serially passaging a bacterial species with no documented history of an association with metazoan hosts, *Shewanella oneidensis* MR-1, through the digestive tracts of a model vertebrate, zebrafish (Figure 1). We chose this *Shewanella* strain because it is one of the best studied bacterial strains isolated from a non-host environment, and it is genetically tractable [30]. We predicted these attributes would maximize our ability to map evolved phenotypes to genotypes.

**Figure 1:**
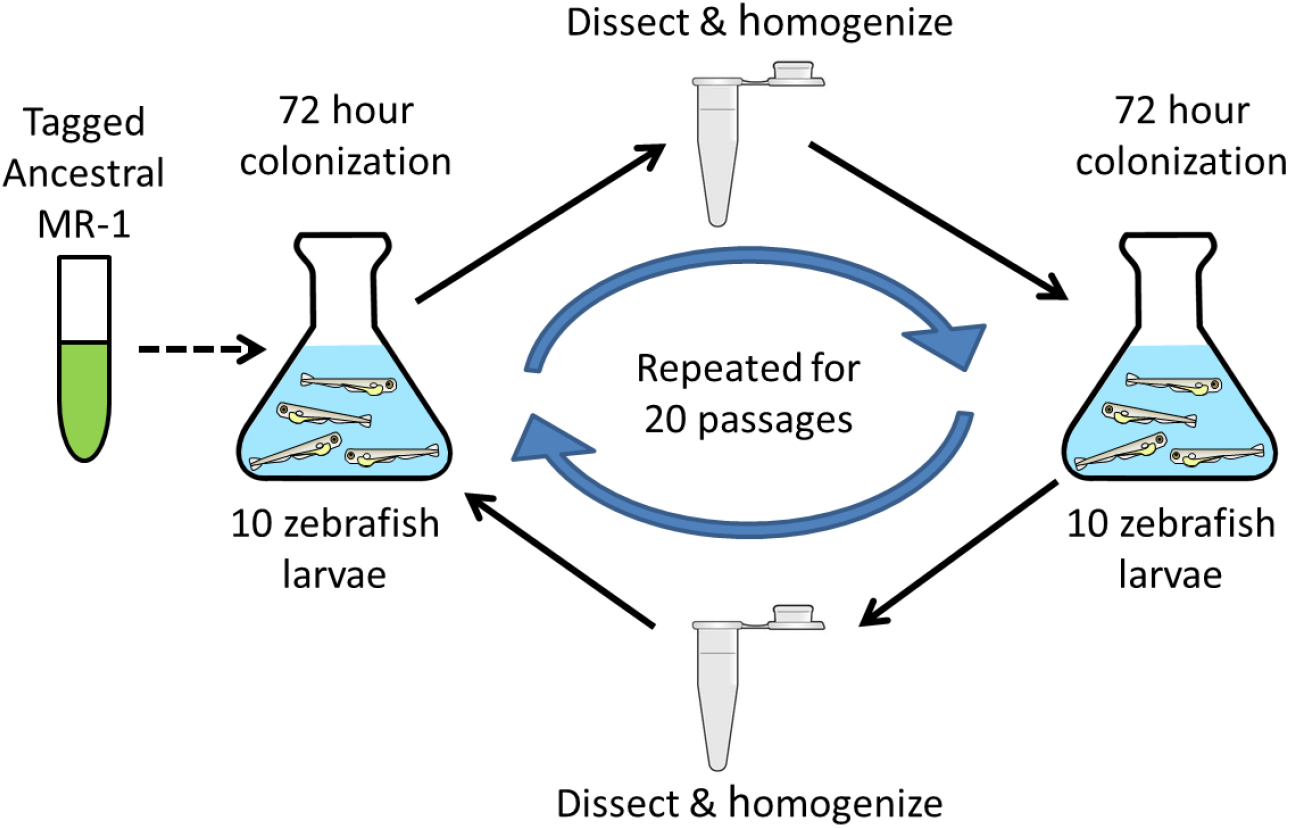
Serial passage schematic. GF larvae were incubated with *MR-1* populations for 72 hours, then 10 larval guts were dissected and homogenized. A sample of the homogenate was used to inoculate a subsequent set of GF larvae. This cycle was repeated for 20 passages. The dotted arrow indicates that no additional *S. oneidensis* ancestor was added to the experimental system after the first passage.

After 20 passages, we observed that evolved isolates from five of the six replicate lines demonstrated a significantly improved ability to colonize larvae compared to their unpassaged ancestor. We then characterized the two evolved isolates with the highest average relative fitness and discovered that each isolate had accumulated a distinct set of mutations. Interestingly, despite these different mutation profiles, both isolates evolved augmented swimming motility relative to the ancestral reference strain, demonstrating phenotypic parallelism in the adaptive trajectories of these two independently evolved isolates. Our findings show that bacteria can rapidly evolve novel host-associations and suggest that swimming motility is advantageous for colonizing aquatic hosts.

## Results

### MR-1 colonizes zebrafish at lower densities than a closely-related Shewanella zebrafish isolate

Gut-associated bacteria are routinely isolated from their animal hosts. Although MR-1 has never been found within a host gut, other strains belonging to the *Shewanella* genus are common in the larval zebrafish microbiota [31]. Indeed MR-1 shares a recent common ancestry with a *Shewanella* species that was isolated from the zebrafish gut (Figure 2; *Shewanella* ZOR0012, Shew-Z12 from this point forth). Interestingly, a whole genome comparison revealed that MR-1 shares an average nucleotide identity (ANI) of approximately 89% with Shew-Z12, as compared to 72% with the more distantly related *Shewanella woodyi* (Figure 2). The high degree of overlap between MR-1 and Shew-Z12 was further reflected when we compared the protein sequence alignments of Shew-Z12 or *S. woodyi* against our MR-1 reference genome (Figure S1). Relative to *S. woodyi*, MR-1 displays much higher levels of amino acid sequence identity with Shew-Z12 on a per gene basis, implying greater amounts of functional conservation between MR-1 and Shew-Z12 (Figure S1).

**Figure 2:**
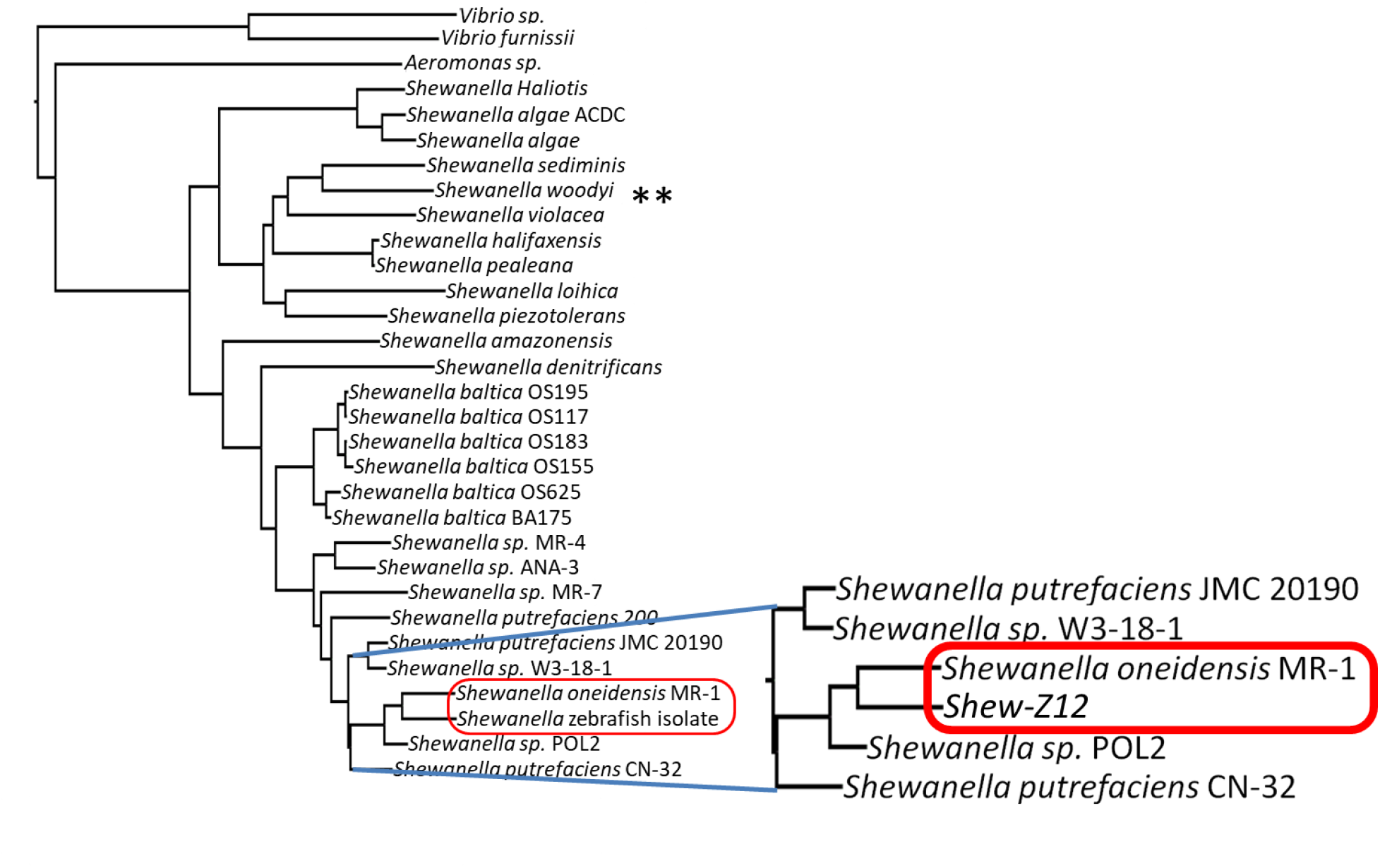
Relatedness of MR-1 to other *Shewanella* species. A phylogenetic tree, based on 16S gene sequences (Table S2), showing the relationship between multiple different *Shewanella* species.

Given the close phylogenetic relationship between MR-1 and Shew-Z12, we wanted to determine whether MR-1 and Shew-Z12 would have similar larval gut colonization characteristics. To assess this, we compared the ability of each strain to colonize GF larval zebrafish guts under both monoassociation and competitive conditions. In both scenarios, flasks containing 10-15 GF larval zebrafish were inoculated with bacterial densities of ~103 colony forming units per mL (CFU/mL). Gut colonization was assessed by dissecting and plating larval digestive tracts after 72 hours of exposure. In monoassociation MR-1 colonized GF larvae at about 10-fold lower densities than Shew-Z12 (Figure 3A). To assess the competitive fitness of these strains, we competed a strain of MR-1 tagged with a neutral fluorescent marker against Shew-Z12 at a one-to-one ratio and quantified relative fitness using a competitive index. The competitive index was calculated by dividing the ratio of each competitor observed in dissected guts by the ratio of each competitor in the inoculum. We found that MR-1 significantly underperformed the Shew-Z12 isolate, and that this difference was greater than could be explained by the difference in abundances in monoassociation (Figure 3B). Given that MR-1 was neither able to colonize larvae to the same capacity as Shew-Z12 in monoassociation, nor able to compete effectively with this closely related host isolate, we concluded that MR-1 has the potential to improve this host association. We speculated that, given its recent divergence from Shew-Z12, adapting MR-1 to the host from which Shew-Z12 was isolated could provide some insight into the adaptive pathways available to bacteria as they transition to life as a host-associated symbiont.

**Figure 3:**
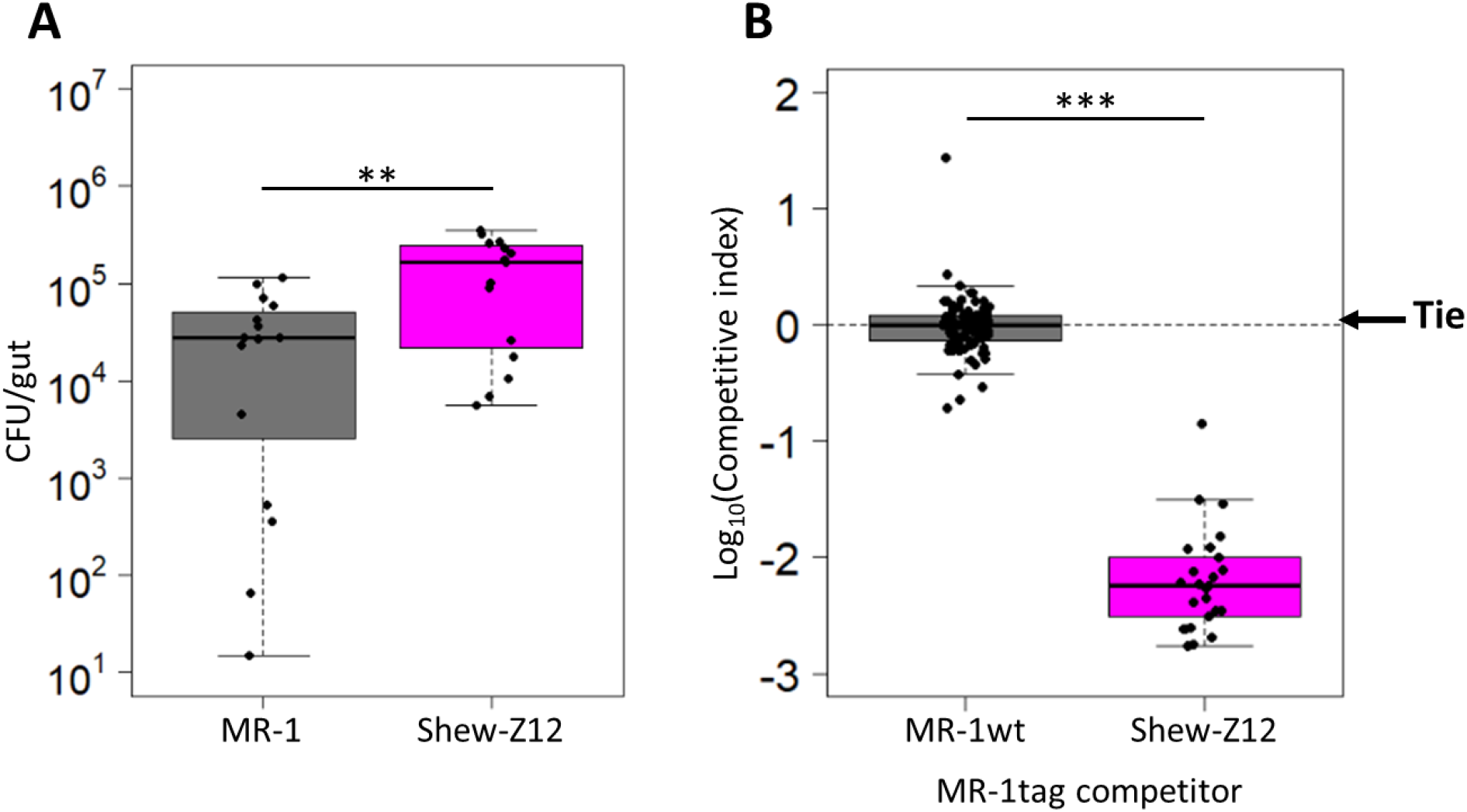
Fitness comparison between MR-1 and Shew-Z12. A) Colonization density achieved in larval guts after 72 hours of colonization under monoassociation conditions for indicated strains. Dissected guts were plated on TSA and colony-forming units (CFUs) were counted. Each point represents a single dissected gut. B) Competitive ability of unpassaged fluorescently-tagged MR-1 strain competed against an untagged version of itself (MR-1wt; left column), or an untagged Shew-Z12 (right column). Each point represents the competitive index measured for a single larval gut (see Methods for details on how competitive indices were calculated).

### Serial passage increased fitness in the gut

To understand how MR-1 would adapt to a vertebrate host gut, we serially passaged six replicate populations through the digestive tracts of GF larval zebrafish (Figure 1A). At the start of the experiment, each population was composed of two fluorescently-marked MR-1 isolates (dTomato, MR-1dT; gfp, MR-1gfp) so that passaged populations could be distinguished from the unpassaged ancestor, and adaptive events could be inferred from changes in each tag’s frequency within the evolving populations (Figure S2). After 20 passages we assayed the fitness of a single, randomly selected isolate per evolved population by competing each isolate against its unpassaged ancestor. Each competition was performed as described above for the MR-1 versus Shew-Z12 competitions. Of the isolates we tested, five of the six outcompeted the ancestral strain, exhibiting competitive indices that were significantly different from the ancestral control group (Figure 4A). These improvements were not likely due to adaptation to the general lab environment, because competitions between replicate evolved populations and the MR-1 ancestor in rich media (tryptic soy broth) produced competitive indices not significantly different from 1.0 (Figure 4B).

**Figure 4:**
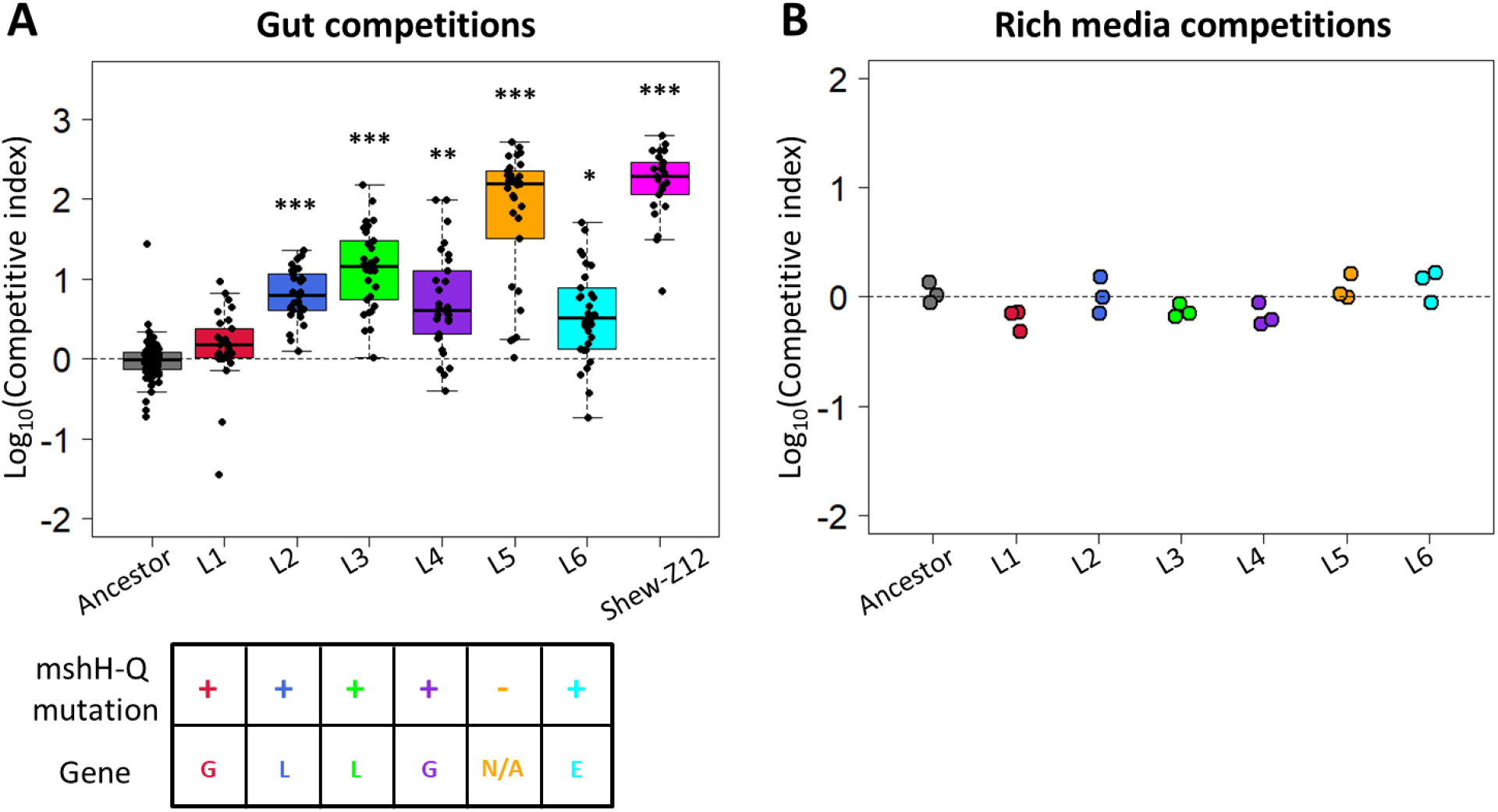
Competitive fitness of evolved *S. oneidensis* isolates. A) Competitive ability of MR-1 isolates from each replicate evolved population against the ancestral MR-1 reference strain. An ancestral competition between tagged and untagged ancestors (left-most box) is shown as a control to show the neutrality of the fluorescent tags. A linear model was used to compare each group against the ancestral control (p < 0.05: *, p < 0.01: **, p < 0.001: ***). Each point represents the competitive index measured for a single larval gut culled from at least 3 replicate flasks (n = 89 guts for ancestor versus ancestor, 25 guts for Shew-Z12 versus ancestor, and 30 guts for all other groups.). The grid at the bottom indicates whether each evolved isolate contained a mutation in the msh operon. Top row: (+) indicates presence of msh operon mutation, (−) indicates absence of msh operon mutation. The bottom row indicates which gene msh operon mutations could be found within. B) Competitive ability of each replicate evolved MR-1 population competing against the ancestral MR-1 reference strain in tryptic soy broth (TSB). Each point represents the competitive index measured for a single biological replicate.

### Comparative genomics revealed multiple candidate adaptive mutations

To investigate the genetic determinants of the adaptation, we sequenced the genomes of each passage 20 evolved isolate. The reads from each isolate were aligned to the ancestral reference genome to identify mutations that have accumulated during serial passage. This analysis revealed numerous mutations per genome (Table S1), which made it difficult to distinguish adaptive mutations from non-adaptive mutations. For example, mutations with neutral or even slightly deleterious impacts on fitness could have increased in frequency simply through their linkage with another, more beneficial mutation.

Therefore, to infer adaptive mutations, we focused on mutations that had accumulated in similar genomic regions across several of our evolved isolates, as such events would be unlikely to occur by chance. In five of the six evolved isolates, we observed mutations in the mannose sensitive hemagglutinin pilus operon (mshH-Q; Figure 4A; Table S1, yellow shaded) suggesting that the msh pilus encoded by this operon likely influences larval gut colonization. These mutations were located in the mshG (L1 and L4), mshL (L2 and L3), and mshE (L6) genes (Figure 4A; Table S1), which encode integral membrane platform, secretin, and ATPase proteins respectively [32]. Interestingly, the msh pilus has been shown to play a role in a number of other host-microbe systems. For instance, within the *Vibrio* genus, msh pili have been suggested to be important for adherence to human intestinal cells [33], as well as colonization of the digestive tract of *Caenorhabditis elegans* [34], and the light organ of *Euprymna tasmanica* [35]. Additionally, in both *Vibrio* and *Pseudomonas* systems, evidence suggests this pilus can interact with components of the mammalian immune system [12, 36]. These observations strengthen the hypothesis that MR-1’s msh pilus was a target of selection in our study.

Notably, the L1 isolate contained a mutation in the mshG gene (similar to L4) but did not have a significant colonization advantage over the ancestor (Table S1). If the other two mutations in L1 were deleterious and recently acquired, it could reconcile L1’s nonadaptive performance with its presence in its population after 20 passages. Alternatively, if the specific mshG mutation observed in L1 and L4 is not advantageous, the difference in relative fitness exhibited by these two isolates might be explained by the additional mutations that are unique to each isolate. More detailed evolutionary genomic analyses that clarify the chronology of accumulated mutations in our evolved isolates could help to establish the basis for L1’s apparent lack of adaptation.

To assess whether the msh operon mutations we observed provided evidence that MR-1 was on a similar adaptive trajectory to one potentially taken by Shew-Z12, we compared the msh operon (mshH-Q) of our reference MR-1 strain to that of Shew-Z12. If evolved MR-1 isolates were on a convergent adaptive trajectory with Shew-Z12, we expected that the mutations found in our evolved isolates would result in amino acid residues similar in identity or biochemistry to those found in Shew-Z12. However, based on our msh operon alignment we did not find any evidence that differences between the MR-1 and Shew-Z12 were ameliorated by evolved mutations. Our alignment did reveal several regions with elevated divergence between ancestral MR-1 and Shew-Z12 (Figure 5). In particular, Shew-Z12’s mshQ gene appeared to be noticeably larger than MR-1’s mshQ gene. The fact that there were substantial regions of divergence between the mshH-Q amino acid sequences of MR-1 and Shew-Z12 left open the possibility that the adaptive changes we observed might amount to evolutionary convergence at the functional or regulatory level. Determining this definitively would require more mechanistic and expression-focused approaches respectively.

**Figure 5:**
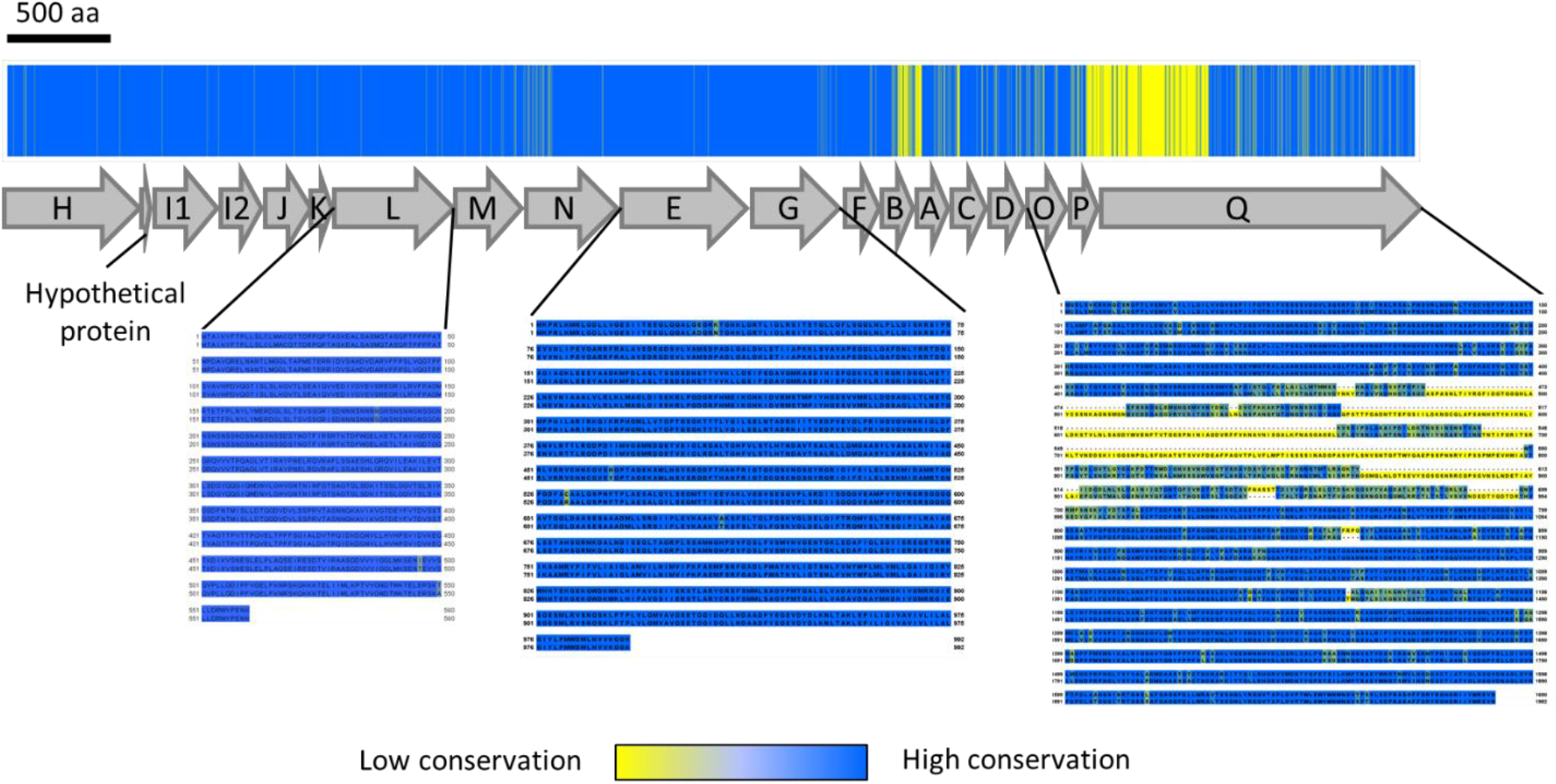
MshOP amino acid conservation between MR-1 and Shew-Z12. The bar across the top shows the mshH-Q amino acid conservation between MR-1 and Shew-Z12. A schematic depicting the organization of the mshH-Q is shown below. Per site, blue indicates the same amino acid is present in both species, while yellow indicates mismatches. Aligned sequences are shown for genes in which we observed mutations in our evolved isolates. In each row of the displayed sequences, MR-1 is featured on top and Shew-Z12 is featured on the bottom. MshQ did not accumulate mutations in our experiment, but it is shown to illustrate that much of the divergence in this protein resulted from a difference in length between MR-1 and Shew-Z12. The color coding in each sequence alignment indicates the degree of conservation. Amino acids with similar biochemistry are bluer, while those with divergent biochemistry are more yellow. A scale bar above the figure indicates the length of 500 amino acids.

The L5 isolate, which exhibited the highest mean fitness of the isolates we tested, contained an entirely unique set of mutations that were not observed in any other evolved isolates. This raises the potential that the L5 isolate may be on a divergent adaptive trajectory, evolving unique phenotypes associated with the exploitation of a distinct niche. Alternatively, it is also possible that L5 may be on a parallel phenotypic evolutionary trajectory, evolving phenotypes similar to those of the other evolved isolates, although via a different set of mutations. To distinguish between these hypotheses, we examined the phenotypes of aggregation behavior and motility in isolate L5 and in isolate L3, the msh-mutant-containing isolate with the highest mean fitness (Figure 4).

Several recent studies have demonstrated that aggregation and motility behaviors can have deterministic impacts on the spatial distribution and competitive dynamics of bacteria within the larval zebrafish gut [37–39], and aggregative behaviors are thought to be a crucial step during successful infection by several human pathogens [33, 40, 41]. Additionally, shifts in motility could impact the rate with which our evolved isolates encounter larval hosts or navigate to their optimal habitat within the gut [14]. Therefore, changes in either of these phenotypes could alter how evolved isolates compete with the ancestral MR-1 strain, and differences in these phenotypes between the L3 and L5 isolates could suggest they have adapted to separate niches within our host system.

### L3 and L5 isolates exhibit different biofilm phenotypes

We compared the capacity of each isolate to form biofilms using static polystyrene plate-based biofilm assays in larvae conditioned medium (LCM). This medium was collected from flasks of GF larval zebrafish at four days post fertilization – the time when we inoculate larvae during competition assays – and it should provide a similar nutrient profile to that experienced by MR-1 during our evolution experiment. We found that L5’s biofilm phenotype was comparable to that of the ancestor, while L3 had a reduced biofilm phenotype (Figure 6). We also assessed how the biofilm phenotypes of the L3 and L5 isolates compared to that of Shew-Z12 and found that Shew-Z12 had a mean biofilm phenotype that was intermediate between L3 and L5, although not significantly different from the MR-1 ancestral strain (Figure 6). Given that L3, L5, and Shew-Z12 are all capable of outcompeting the MR-1 ancestor in their ability to colonize GF larval zebrafish guts, yet these strains display a range of biofilm phenotypes under the conditions tested, we conclude that our biofilm assay is not capable of predicting competitive fitness *in vivo*. However, our results demonstrate that the unique mutational profiles of the L3 and L5 isolates result in distinct aggregative behaviors.

**Figure 6:**
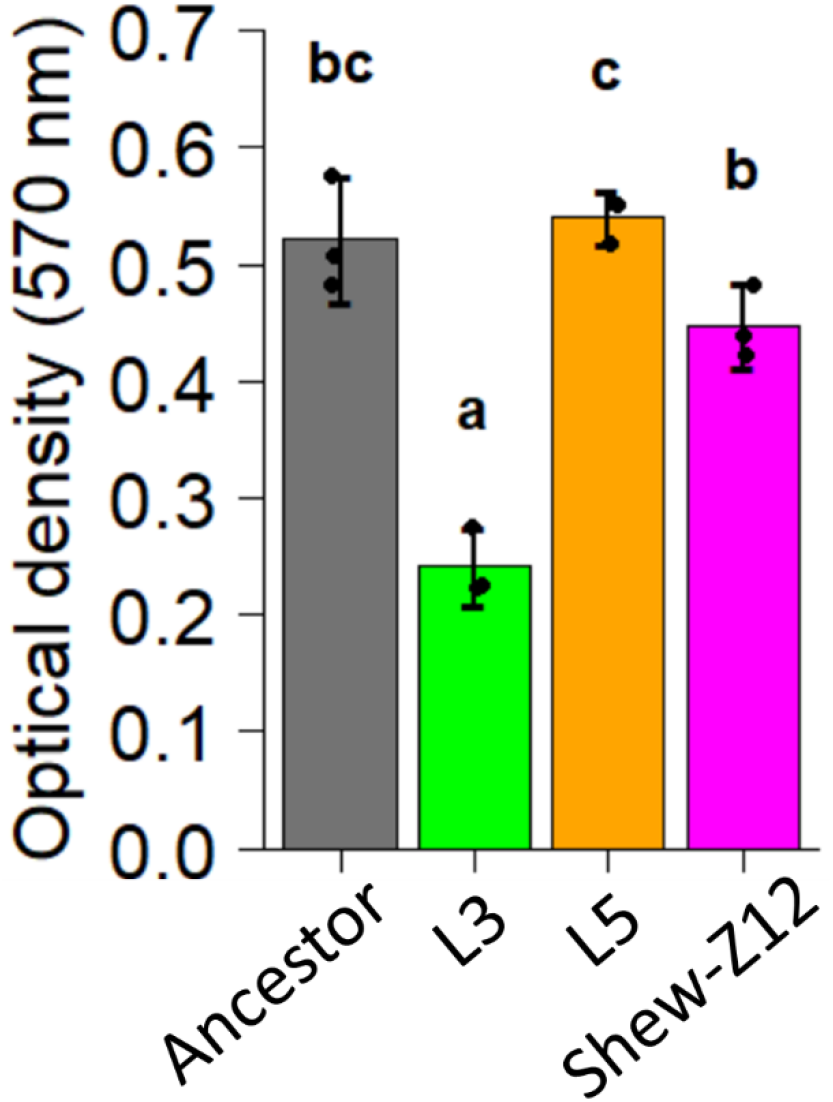
Crystal violet biofilm assay comparing L3, L5, and *Shewanella sp. ZOR0012* to ancestor. Optical density (570 nm) corresponds to crystal violet intensity. Higher optical density readings indicate more robust biofilms. Statistical groupings are indicated by letters above each bar for a significance threshold of p<0.05. Letters in common between groups indicate the absence of a significant difference in each group’s mean. Only the L3 isolate shows a significantly altered biofilm phenotype compared to the wt ancestor.

### L3 and L5 are more motile than the ancestor

Since Robinson and colleagues previously demonstrated that serially passaged *Aeromonas* populations, which were more motile than an unpassaged ancestor, were able to better colonize GF larval zebrafish [29], we hypothesized that MR-1 could similarly improve host colonization in our study via enhanced motility. A common assumption in microbiology is that there is a trade-off between adherence and motility [42–45], wherein cells that tend to be more adherent also tend to be less motile and vice versa. Given the difference we observed between the ability of L3 and L5 to adhere to surfaces, we wondered whether these strains might also exhibit a difference in motility.

We quantified the cellular swimming speeds and motile population fractions of these strains within flasks of GF larvae, by analyzing fluorescence microscopy images with an automated cell tracking algorithm. These assays were conducted in competition (evolved isolate versus ancestor) to mimic the conditions under which we assessed fitness. Thus, if differences in motility were dependent on the competitive dynamics between the evolved isolates and their ancestor, we would capture that in these assays. Surprisingly, despite their differing biofilm phenotypes, we found that both the L3 and L5 isolates were more motile than the ancestor. The evolved strains demonstrated both faster speeds (Figure 7, A and B), and they had a larger portion of their population that was motile (Figure 7C). These results confirm that motility can be important for larval zebrafish colonization.

**Figure 7:**
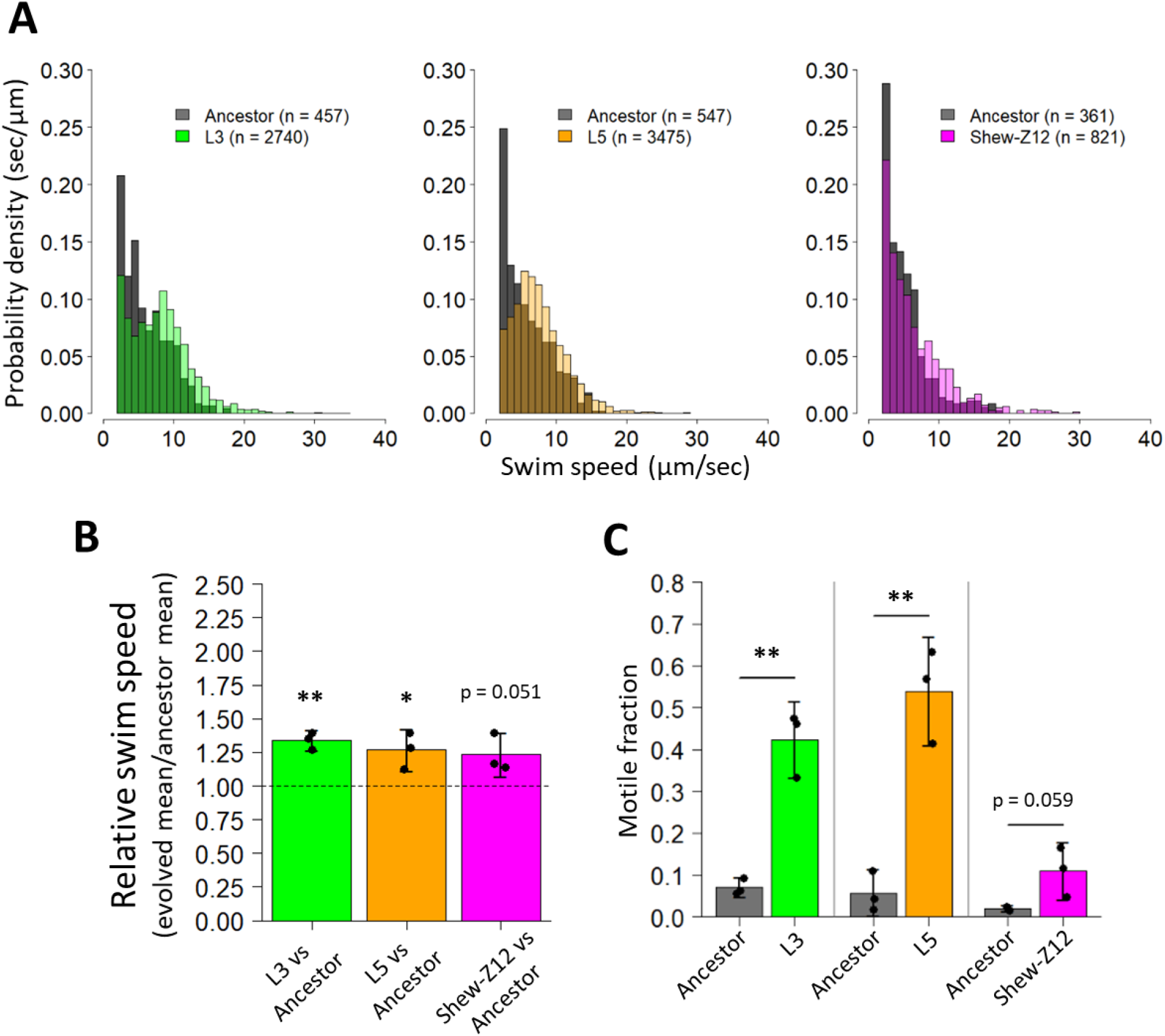
Motility characteristics of evolved and zebrafish isolates compared to ancestor. A) Histograms comparing the swim speeds of L3, L5, and Shew-Z12 with the ancestral MR-1 strain are shown based on aggregated observations from three flasks per comparison. B) Mean swim speed of L3, L5, and Shew-Z12 relative to the ancestor. For each flask, represented by a point, the mean mutant swim speed was divided by the mean ancestral swim speed. One-tail t-tests against a mu value of one (our null expectation) were conducted to assess whether each group exhibited swim speeds that were significantly faster than the ancestor (p < 0.05: *, p < 0.01: **). C) The fraction of L3, L5, Shew-Z12, and ancestral MR-1 populations that were observed to be motile (swimming speeds > 2 μm/sec). Each point represents the motile fraction for a single flask, and each bar represents data from 3 separate flasks. For each panel, the L3 and L5 comparisons were done under competition, while the Shew-Z12 comparison was done under monoassociation conditions. Error bars indicate the 95 % confidence interval.

We next compared the motility phenotypes of our MR-1 ancestor and evolved isolates to the zebrafish isolate. Our hypothesis was that Shew-Z12 would also exhibit greater motility than the MR-1 ancestor. We examined Shew-Z12 motility as described above, with the exception that Shew-Z12 and our ancestral MR-1 strains were assayed in monoassociation. Attempts to fluorescently tag Shew-Z12 were unsuccessful, leaving no way to distinguish this species via fluorescence microscopy, and thus we collected data for these assays in bright field. We found that Shew-Z12 too exhibited faster swimming speeds and a greater motile fraction of the populations compared to the MR-1 ancestor (Figure 7). This supports the hypothesis that in our study MR-1 evolved along an adaptive trajectory that is phenotypically convergent with Shew-Z12.

## Discussion

We serially passaged replicate bacterial populations through the intestines of GF larval zebrafish to explore how a non-host-associated bacterium adapts to a novel vertebrate host. We expected that there would be an abundance of niche space available to evolving MR-1 populations. For example, cells capable of colonizing larval guts must compete externally in the aqueous environment in order to access the larvae, and then compete *in vivo* so they will be sampled and carried over in subsequent passages. Both environments likely contain unique sets of selective pressures. On one hand, we hypothesized that this might select for divergent adaptive trajectories resulting in unique genotypes, and associated phenotypes, that would allow for the exploitation of distinct niches. Alternatively, if the selection resulted in adaptation to a single niche within our system, we expected that specialists for that niche would exhibit high relative fitness, leading to phenotypic parallelism among adaptive lineages. Such parallelism could indicate that only a small number of traits are likely to facilitate the initiation of host associations. Moreover, because traits that are common among independent adaptive genotypes have an increased likelihood of playing a causal role in enhancing fitness, such parallelism could provide evidence that phenotypes are adaptive. To address these hypotheses, we examined the mutations and phenotypes of two evolved isolates with the highest mean fitness in our experimental system: isolates from the L3 and L5 populations.

Consistent with expectations from past experimental evolution studies [19], we observed that independently evolving populations of MR-1 rapidly produced adaptive isolates after serial passage through larval zebrafish. Interestingly, the improved competitive fitness we observed in the L3 and L5 isolates was associated with unique genotypes (Table S1), and this suggested that each isolate found idiosyncratic strategies for adaptation. L3 contained a missense mutation in the mshL gene, and mutations in the msh operon were observed in five out of six of the evolved isolates we sequenced (Figure 4A, lower grid; Table S1), which supports the conclusion that these mutations were likely adaptive. The msh pilus has also been implicated in several other host-microbe systems [12, 33–36], and therefore focusing on the products encoded by this operon may yield fruitful avenues for researchers interested in manipulating host-microbe interactions.

Less clear is which mutations in the L5 isolate’s genome are adaptive. The mutations we identified fell within genes that were annotated as a sensor histidine kinase, a lipoprotein, a diguanylate cyclase, and a hypothetical protein (Table S1). These mutations could affect a broad range of cellular and physiological processes, however without further investigation, it is difficult to predict which of these mutations actually plays an adaptive role in MR-1’s ability to colonize larvae. We plan to disentangle the effect of these mutations in future work.

With respect to the biofilm phenotypes we assessed, the L3 isolate displayed a reduced ability to form biofilms, while neither the L5 isolate nor Shew-Z12 exhibited biofilms that differed significantly from the MR-1 ancestor phenotype. Although this suggests biofilm formation does not correlate with competitive fitness, the cell densities required to generate detectable biofilms in these assays are orders of magnitude higher than would be experienced by MR-1 in a larval flask. Therefore, it is possible the aggregative behaviors we observed are not representative of the behaviors MR-1 isolates would manifest when competing to colonize the larval gut. Nonetheless, the ability to form robust biofilms could pose several advantages with respect to host colonization. Bacteria are known to form intimate associations with host epithelial tissues, and the ability to form biofilms can be critical to successful host colonization, and host specificity [46, 47]. Additionally, bacterial aggregates can shield internal members from harsh environmental conditions until more favorable conditions are encountered within a host, wherein large numbers of cells can detach and colonize [48]. Similarly, upon colonization of a host, biofilms can also protect bacterial members from harmful host defenses [49]. Given these potential advantages, it makes sense that the evolved isolate with the highest mean fitness, L5, would maintain a robust ability to form biofilms.

Conversely, the L3 isolate’s reduced biofilm phenotype may be explained by the mshL missense mutation it contained. MshL encodes a putative outer membrane pore protein through which the msh pilus extends [32, 50–52]. Consistent with what others have found for the msh pilus, if this mutation was a loss of function mutation, it could reduce MR-1’s msh pilus expression – and its ability to form biofilms – resulting in a more planktonic existence [52–55]. In turn, this could increase L3’s encounter rate with zebrafish larvae that are swimming through the aqueous environment. In this way, the L3 isolate would have a competitive advantage over ancestral cells that were adherent to flask surfaces and were therefore less able to access larval hosts. Alternatively, reduced expression of the msh pilus could potentially help the L3 isolate evade elements of the larval immune system upon gut colonization [12, 36]. Neither of these hypotheses necessarily accounts for the L3 isolate’s faster swimming speeds, although speculatively, it is possible that if MR-1 were not producing pili, it might be able to devote more resources to motility which is energetically costly [56].

In our study, the fact that the L3 and L5 isolates evolved to outcompete the ancestor, while exhibiting distinct biofilm phenotypes, suggests that each isolate could be pursuing alternative adaptive strategies. However, even though biofilm formation often trades off with motility [43–45], we observed selection for enhanced motility via two separate genetic pathways, which lends support to the conclusion that augmented motility is adaptive. Further, this phenotype was also observed in the closely related zebrafish isolate, Shew-Z12. Interestingly, the motile fraction of Shew-Z12’s population is reduced compared to either the L3 or L5 isolates (Figure 7C). This could stem from the fact that Shew-Z12 evolved its motility characteristics in the presence of a bacterial community, whereas our experiment was conducted under germ-free (i.e. axenic) conditions. It is possible that this historical difference may have altered the costs or benefits of motility, leading Shew-Z12 to evolve distinct motility characteristics. For example, cross-feeding between community members could alter bacterial foraging requirements and result in unique motility optimization [57, 58].

Although it is not clear exactly how motility improves host colonization in our system, one possibility is that enhanced motility increases chemotactic responses to host-produced chemical gradients, thereby increasing bacterial encounter rates with larval zebrafish. Alternatively, once an evolved MR-1 cell encounters a host, faster swimming speeds could help strains traverse narrow junctions within the host to reach the gut more quickly than their ancestral competitor [14]. However, prior work showed no fitness advantage for a hypermotile *Aeromonas* strain gavaged into the oral cavity of larval zebrafish, suggesting this mode of fitness enhancement may be less likely [29]. Another option is that after bacterial cells migrate into the digestive tract, motility could help bacteria resist expulsion induced by intestinal contractions [39]. Ultimately, our results add to a growing body of evidence implicating motility as an important trait for host colonization [14, 17, 29, 59], and the phenotypic parallelism we observed suggests that traits associated with dispersal can play a critical role in the establishment of host-microbe symbioses [60].

## Acknowledgements

This research was supported by the National Institutes of Health (grants T32GM007413, T32GM007759, and P01GM125576). Additionally, we would like to thank the University of Oregon Zebrafish Facility and Karen Guillemin for their support and resources. We would also like to thank Tristan Ursell for aiding in fluorescence microscopy image acquisition and Daniel Shoup for fluorescence microscopy training; Raghuveer Parthasarathy for providing cell-tracking image analysis software. Finally, we would like to thank the personnel of the Bohannan, Guillemin, and Parthasarathy laboratories for their intellectual contributions, with a special thanks to Beth Miller, Hannah Tavalire, Elena Wall, and Travis Wiles.

## Methods

### Zebrafish husbandry

To ensure animal specimens were treated ethically in all experiments involving zebrafish, we adhered to the standard protocols and procedures approved by the University of Oregon Institutional Animal Care and Use Committee (IACUC protocol: 15-98). GF derivations were carried out as described in Melancon et al., 2017. Details about larval gut dissections can be found the Serial passage section.

### Bacterial strains

Our ancestral reference *S. oneidensis* (MR-1) and Shew-Z12 strains were obtained from Karen Guillemin’s laboratory at the University of Oregon. GFP and dTomato tags were inserted using the Tn7-based approach described in Wiles et al., 2018. All *S. oneidensis* strains were cultured in TSB at 30 °C under shaking conditions.

### Serial passage

Our experimental evolution serial passage scheme was similar to the one outlined by Robinson et al. 2018. 5 mL overnight tryptic soy broth (TSB) cultures of MR-1 isolates tagged with either green fluorescent protein (MR-1gfp) or dTomato fluorescent protein (MR-1dT) were diluted 1:100 in TSB and allowed to grow out to late log phase (4-5 hours). Six replicate ancestral populations were then generated by combining subcultures of MR-1gfp and MR-1dT at a 1:1 ratio. These mixtures allowed us to infer the occurrence of adaptive events based on fluorescent tag frequency changes observed throughout the experiment. Beneficial mutations occurring in a tagged genomic background should cause the frequency of that tag to increase over time. Next, 10 μL of each of these replicate ancestral populations were used to inoculate larval flasks containing ~15 mL of EM and ~15 GF larval zebrafish at 4 days post fertilization (dpf)(inoculating MR-1 densities were ~10^6^ CFU/mL). Larvae were then incubated with MR-1 populations at 28 °C for 72 hours. At 7 dpf, 10 larvae were euthanized with tricane, mounted on a glass slide, and their digestive tracts were dissected out. Glass slides were coated with 3 % methylcellulose to help immobilize larvae during dissections. Following dissections, all 10 digestive tracts culled from each flask were placed in a single 1.7 mL tube containing 500 μL EM and ~100 μL 0.5 mm zirconium oxide beads (Next Advance, Averill Park, NY, US). The contents the larval guts in each of these tubes were then immediately homogenized using a bullet blender tissue homogenizer (Next Advance, Averill Park, NY, US) for 60 seconds at power 4. To preserve our ability to revive replicate populations after each passage, we created freezer stocks by using a pipette to mix 200 μL from each homogenized tube with 200 μL of 50 % glycerol (25% glycerol final concentration). These freezer stocks were stored at −80 °C. The remaining contents of each homogenized tube was then stored at 4 °C for 0-14 days, at which point ~250 μL were sampled to inoculate a subsequent set of GF larval flasks (~15 larvae in ~15 mL EM). Upon inoculation, 100 μL of each larval flask was dilution-plated in triplicate to quantify the inoculating population densities (typically around 10^3^ CFU/mL) and determine tag frequencies. This cycle was repeated for 20 passages. All six replicate evolving populations were maintained separately throughout our experiment.

### Comparative genomics

We submitted our MR-1 and Shew-Z12 strains to the Washington State University Molecular Biology and Genomics Core (WSUGC) for long read sequencing. Genome assembly for MR-1 was conducted by WSUGC, whereas genome assembly for Shew-Z12 was conducted in house with Canu v. 1.7.1 [61]. To generate annotation files for these genomes, we relied on Prokka v1.12 [62], and RAST v. 2.0 [63].

### Phylogenetics

Using Integrated Microbial Genomes and Microbiomes [64] (IMG/M, https://img.jgi.doe.gov/), we collated a set of 16S ribosomal RNA genes from 28 *Shewanella* species, 2 *Vibrio* species, and 1 *Aeromonas* species (See Supplementary Table S2 for metadata). These 16S rRNA genes were entered into Clustal Omega (https://www.ebi.ac.uk/Tools/msa/clustalo/) to generate a multiple sequence alignment file and a subsequent Newick-formatted phylogenetic tree file. This file was then visualized with FigTree v1.4.4 [65].

### Genome comparisons between *S. oneidensis* and other *Shewanella* species

Average sequence identity (ANI) was calculated using the EZBioCloud online ANI calculator [66] to quantify the ANI between *S. oneidensis* and Shew-Z12.

### Specific gene and operon comparisons between *S. oneidensis* and Shew-Z12

ANI was calculated using the same tool described in our whole genome comparisons above. For mshH-Q comparisons, we separately concatenated the amino acid sequences of each gene in the mshH-Q of *S. oneidensis* and Shew-Z12 in series by relying on our RAST-annotated files. we then generated multiple sequence alignment (msa) files using Clustal Omega web tool [67] that compared these mshH-Q sequences and used them to depict sites of divergence along the mshH-Q (Figure 5). Visualizations of single gene or multi-gene comparisons between *S. oneidensis* and Shew-Z12 were created using Clustal Omega-based msa files that were imported into Jalview2 [68]. To highlight regions of divergence within genes we color-coded our comparisons using the color by annotation feature of Jalview2 (Figure 5; Figure 17). This feature color-codes amino acid comparisons per site based on biochemical conservation.

### Evolved mutation calling

We selected one randomly-chosen isolate per evolved replicate population (6 isolates total) by using an inoculation loop to dilution streak a sample from each population’s freezer stock on TSA (tryptic soy agar) plates (one plate per evolved population, totaling 6 plates) and incubating plates at 30 °C for 24 hours. For each population, we overnight cultured four colonies that resulted after 24 hours of growth, and then created freezer stocks (stored at −80 °C) that consisted of a 1:1 mixture of each cultured isolate and an equal volume of 50 % glycerol (final concentration: 25 % glycerol). To create genomic libraries for each evolved isolate, we used an inoculation loop to generate overnight cultures from each isolate’s corresponding freezer stock, and then extracted genomic DNA from each culture using the Promega Wizard® Genomic DNA Purification Kit (Catalog #: A1120). Single-end 150 bp libraries were generated from these genomic DNA extractions according to the Nextera XT DNA Library Prep Kit Reference Guide (Document #: 15031942 v02), and these libraries were sequenced on the Illumina HiSeq 4000. Following this same protocol, we also sequenced the genomes of our ancestral MR-1gfp and MR-1dT strains on the Illumina HiSeq 4000. To identify candidate adaptive mutations, we used breseq 0.31.0 in consensus mode to compare each Illumina sequenced isolate to the annotated ancestral reference separately, and then looked for single nucleotide polymorphisms (SNPs) and indels that were present in each evolved isolate, but absent in MR-1gfp and MR-1dT isolates [69]. With the exception of mshH-Q genes, the gene annotations for the mutations listed in Table S1 were determined by Prokka v1.12 [62]. The mshH-Q gene annotations were determined by RAST v 2.0.

### Competition assays

5 mL overnight TSB (tryptic soy broth) cultures of competing strains were diluted 1:100 in TSB and allowed to grow out to late log stage (4-5 hours). 500 μL of each competitor was then mixed in a single 1.7 mL tube so that competitors were at an approximate 1:1 ratio. Competition mixtures were pelleted (7000 rcf for 5 min) and resuspended in 1 mL sterile EM. Resuspended competition mixtures were diluted 1:100, and 7.5 μL of these dilutions were used to inoculate GF larval flasks containing ~15 mL of EM and ~15 larvae at 4 dpf. Immediately following inoculation, triplicate 100 μL samples from each competition flask were dilution plated to establish the inoculation ratio of competitor1:competitor2 (CFU/mL). After inoculation, flasks were incubated at 28 °C for 72 hours. At 7 dpf, multiple larvae per flask were euthanized with tricane, and their guts were dissected and individually placed in 1.7 mL tubes containing 500 μL of sterile EM and ~100 μL 0.5 mm zirconium oxide beads (Next Advance, Averill Park, NY, US). The contents the larval guts in each these tubes were then immediately homogenized using a bullet blender tissue homogenizer (Next Advance, Averill Park, NY, US) for 60 seconds at power 4. Homogenized tubes were then dilution plated to determine the CFU/gut for each competitor. A competitive index was calculated for each dissected gut by dividing the ratio of competitor1:competitor2 found in each gut by the mean inoculation ratio determined from the triplicate measurements for each corresponding flask, 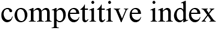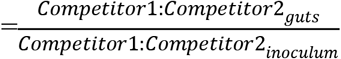. We used Dunnett contrasts to compare dissected larval guts grouped from separate flasks with a linear model where flasks were considered a random effect.

### Rich media competitions

Overnight TSB cultures of competing strains were diluted 1:100 in TSB and allowed to grow out to late log stage (4-5 hours). 500 μL of each competitor was then mixed in a single 1.7 mL tube so that competitors were at an approximate 1:1 ratio. Competition mixtures were pelleted (7000 rcf for 5 min) and resuspended in 1 mL TSB. Resuspended competition mixtures were diluted 1:100, and 5 μL of these dilutions were added to 10 mL TSB in a 20 mL test tube. Immediately following inoculation, triplicate 100 μL samples from each competition culture tube were dilution plated to establish the inoculation ratio of competitor1:competitor2 (CFU/mL). we incubated each competition at 30 °C for 24 hours with agitation (angled back and forth rocker, 60 rpm), at which point we again took triplicate 100 μL samples from each competition tube and dilution plated them to quantify the CFU/mL for each competitor. Competitive indices were calculated by dividing the final CFU ratio of competitor1:competitor2 by the inoculation ratio, 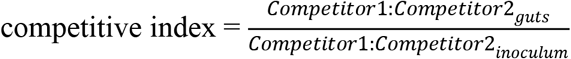

### Biofilm assay

Triplicate biological replicate overnight TSB cultures of strains of interest were diluted 1:100 in TSB and allowed to grow out to late log stage (3-4 hours). 1 mL of subcultured strains were pelleted (7000 rcf for 5 min), and the pellets were normalized to an optical density of 1.0 (600nm) via resuspension with sterile EM. 150 μL of each resuspension were added to four wells of a round-bottom 96 well polystyrene plate (Greiner Bio-One, catalogue #: 650185) per biological replicate. The plate was incubated at 30 °C for 24 hours, and the volume of each well was removed with a multichannel pipette. The wells were rinsed twice with 200 μL of sterile EM, and each well was stained with 180 μL 0.1 % crystal violet (CV). The plate was incubated at room temperature for 10 min, at which point the crystal violet was removed with a multichannel pipette, and the wells were again rinsed twice with 200 μL of sterile EM. The CV was solubilized with 100 % dimethyl sulfoxide (DMSO) for 15 min with agitation (~180 rpm on a rotating minishaker), and 100 μL of the solubilized CV was added to 100 μL of 100 % DMSO in a flat-bottom 96 well polystyrene plate (Corning Incorporated, reference #: 3595). Optical densities were then measured for each well at 570 nm.

### Motility assays

5 mL overnight TSB cultures of strains of interest were diluted 1:100 in TSB and allowed to grow out to late log stage (4-5 hours). Strains were then prepped for inoculation as described above. A volume of 7.5 μL of each strain prepared for inoculation was used to inoculate GF larval flasks containing ~15 mL of EM and ~15 larvae at 4 dpf. Inoculated larval flasks were incubated at 28 °C for 13-17 hours, at which point bacteria in each flask were imaged on an inverted microscope (Nikon Eclipse Ti-e) by focusing on the bottom interior surface of the flask. Ten, 30-45 second movies were then captured at a rate of 15-24 frames per second. When movies were taken of competing populations, each population was fluorescently tagged with either gfp or dTomato, and movies were taken separately to capture the motility dynamics of each tagged population independently within the same flask. Single strain movies were taken in bright field. Bacteria tracking was performed in MATLAB using previously described software (https://pages.uoregon.edu/raghu/particle_tracking.html). In brief, bacteria-like objects were identified by intensity thresholding after bandpass filtering and then localized using a radial symmetry algorithm [70]. Tracks were assembled using nearest-neighbor linking. Tracks shorter than 5 frames were discarded. As an additional filtering step to remove multiple tracks assigned to the same, non-motile bacterium, tracks with a difference in mean position less than 1.7 μm were culled, keeping only the longest track. Based on the measured distribution of swimming speeds, a cutoff of 2 microns/second was used to operationally define a “motile” bacterium from a “non-motile” one.

## SUPPLEMENTS

**Figure S1:**
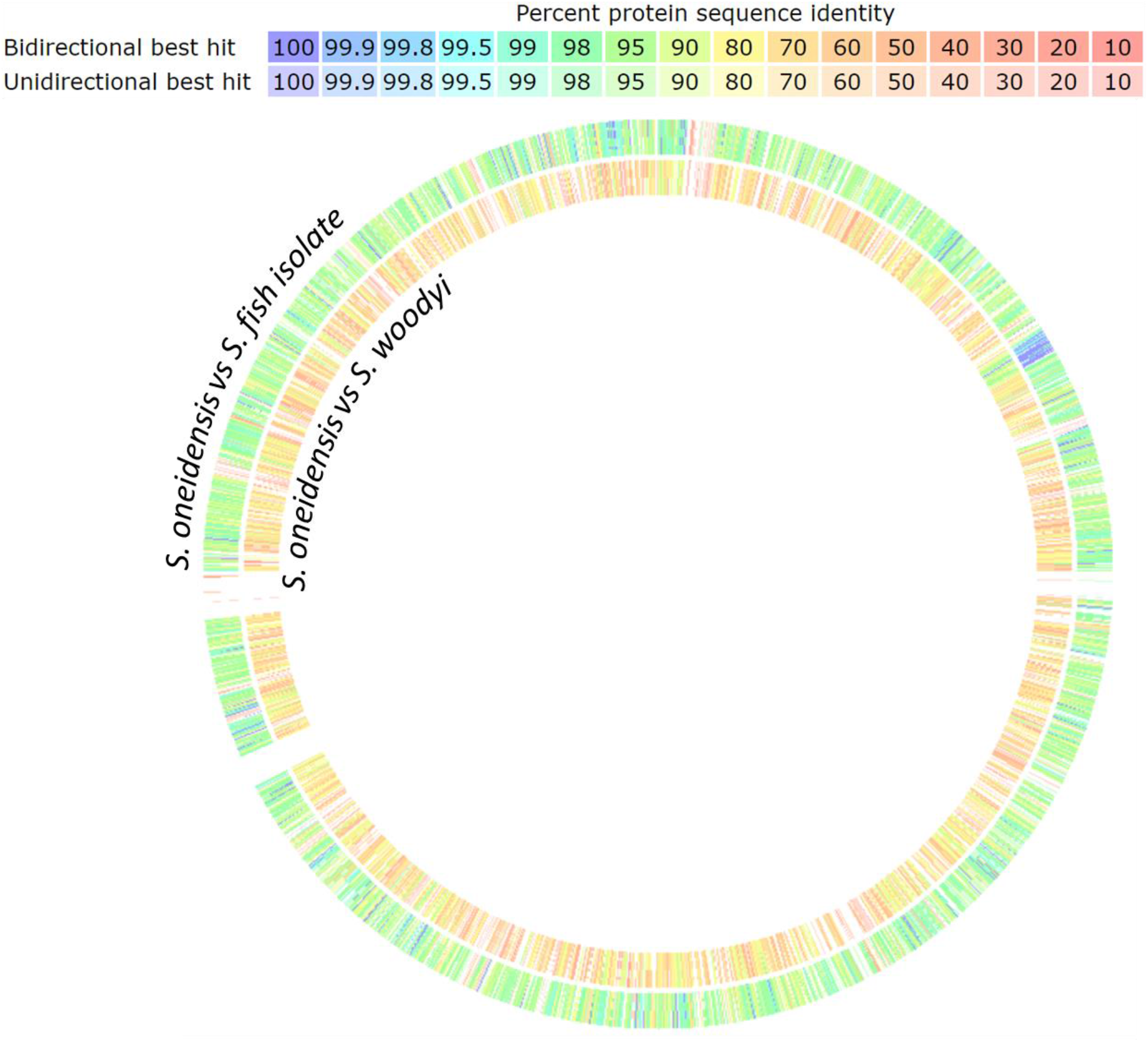
Whole genome amino acid comparisons per gene. Whole genome alignments of Shew-Z12 (outer ring) or *Shewanella woodyi* (inner ring) amino acid sequences against the MR-1 reference genome are shown on a per gene basis. Color shading around each ring indicates sequence identity (key at the top of the figure). Alignments were conducted using the sequenced-based comparison tool of The SEED Viewer v. 2.0 [71].

**Figure S2:**
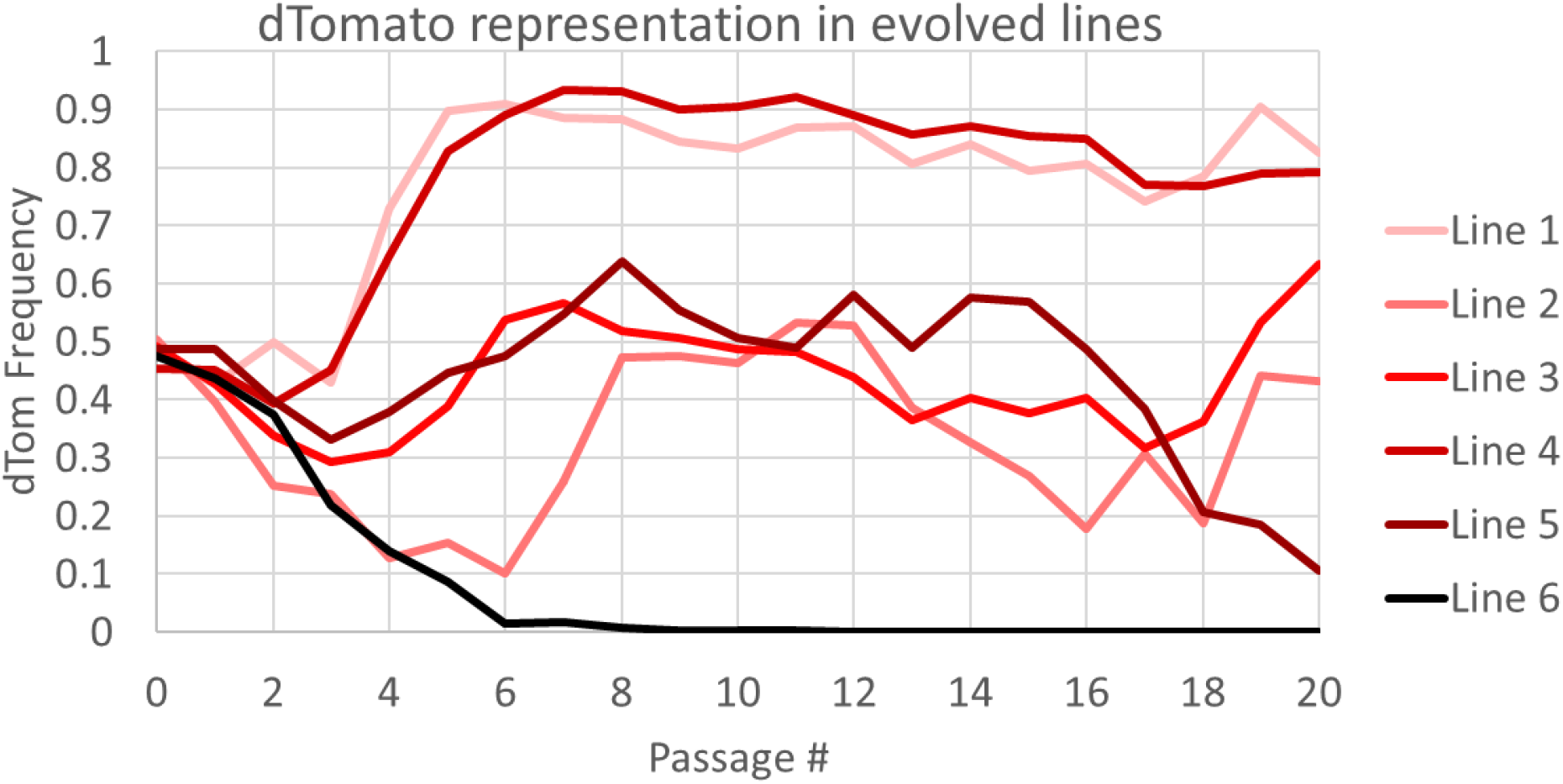
Representation of the dTomato fluorescent tag during serial passage. Shown is the proportion of dTmato-tagged MR-1 CFUs counted for each serial passaged population after each passage.

**Table S1:**
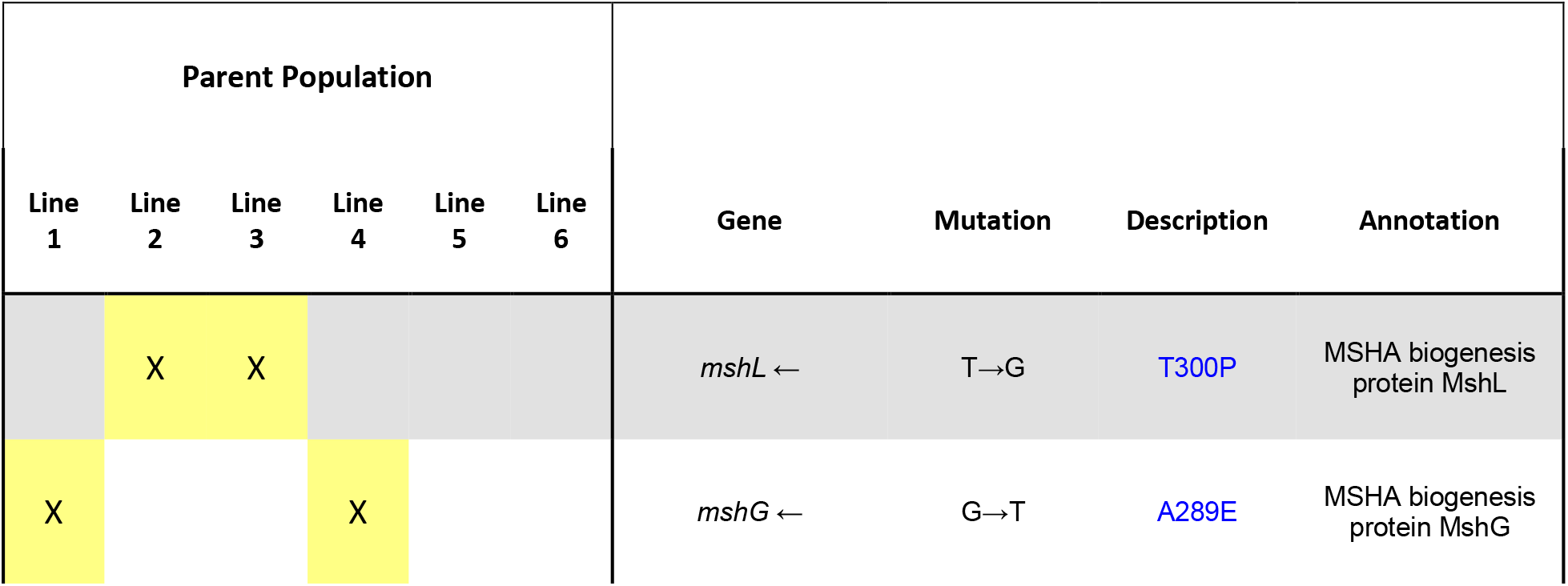
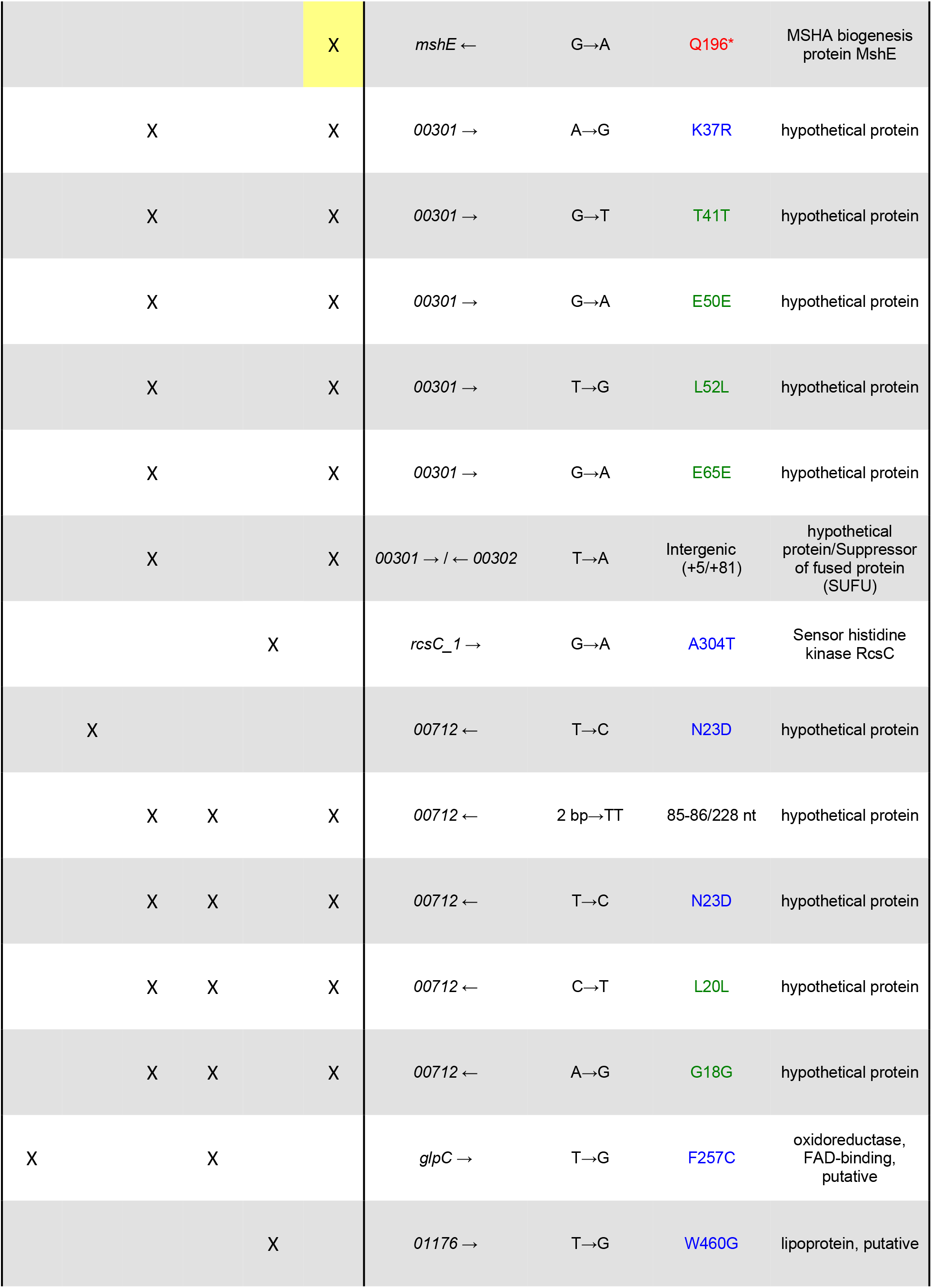
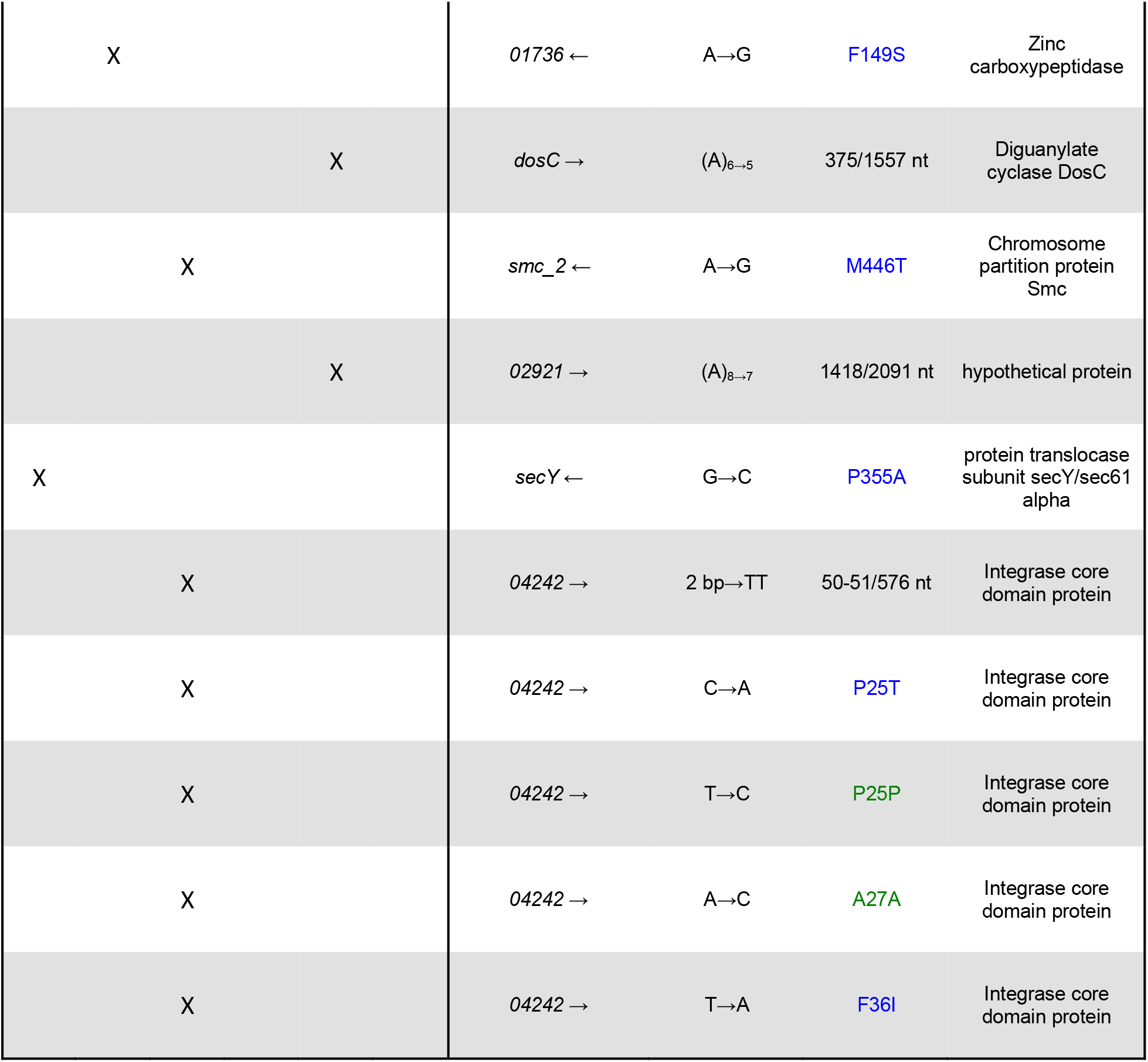
Candidate adaptive mutations. Mutations are listed for four isolates per evolved population. Each isolate is color-coded by the fluorescent tag it contained (red = dTomato, green = gfp). Except for mshH-Q annotations (annotated with RAST v. 2.0), gene annotations are based on Prokka v1.12 (Seemann, 2014). Color-coding for gene descriptions is as follows: green: synonymous mutation, blue: missense mutation, red: nonsense mutation, black: all other mutations. Yellow-shaded cells indicate mshH-Q mutations. Total number of mutations: L1: 3, L2: 3, L3: 17, L4: 6, L5: 4, L6: 11.

**Table S2:**
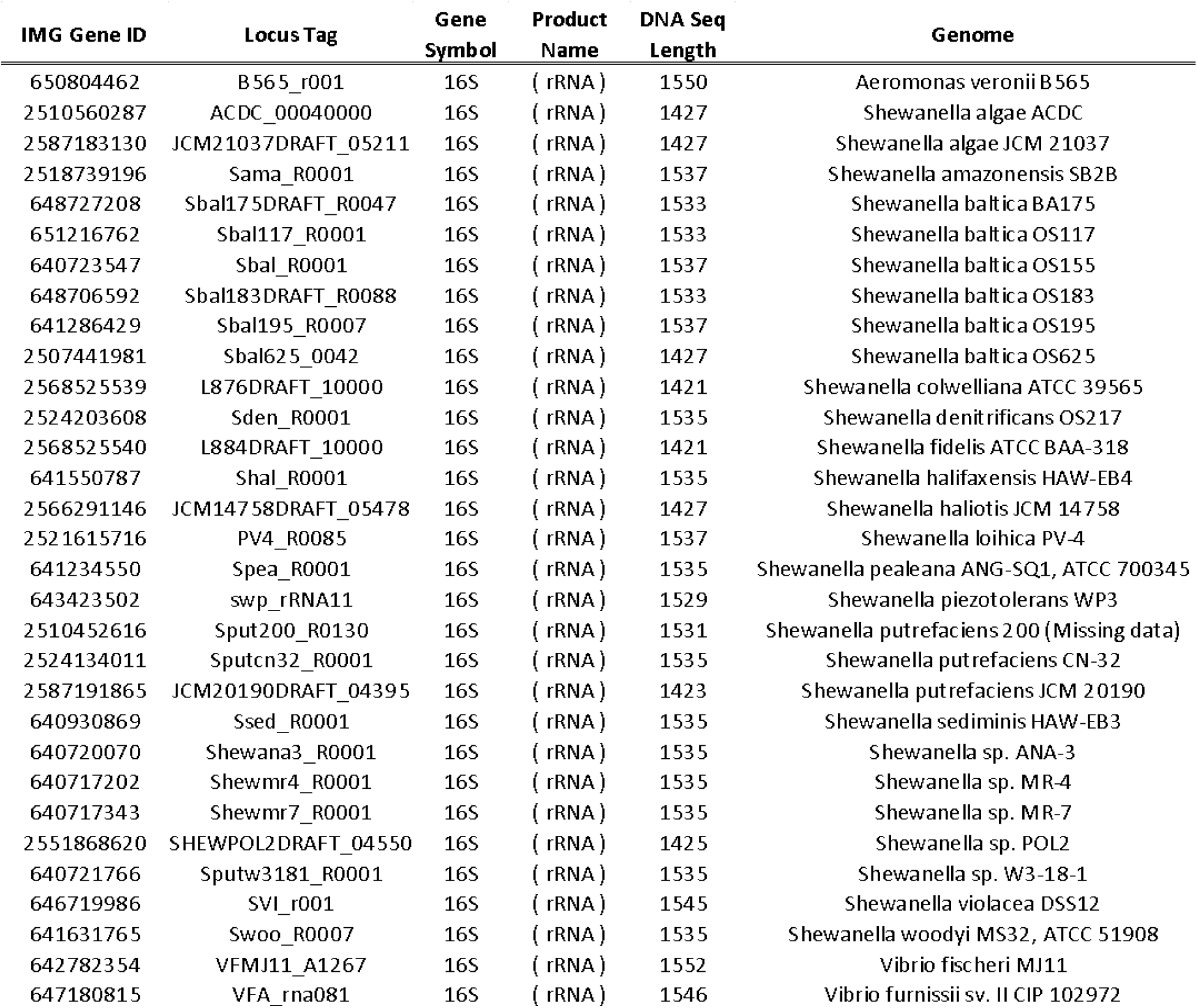
Metadata for 16S genes used to create the phylogenetic tree featured in Figure 2

